# Cryo-EM Structure of Photosystem I from mangroves in their natural environment

**DOI:** 10.1101/2025.06.10.658896

**Authors:** Tirupathi Malavath, Emmanuel Heilmann, Quentin Charras, Xukun Yuan, Ashraf Al-Amoudi, Andreas Naschberger, Azam Gholami

## Abstract

Photosystem I (PSI) is a multi-subunit pigment–protein supercomplex responsible for light driven electron transport in photosynthesis. Its structural plasticity under natural environmental stress remains poorly understood. Here, we present a 2.1 Å cryo-EM structure of the PSI–LHCI supercomplex from *Avicennia marina*, a halotolerant, dominant mangrove species native to the Arabian Gulf that thrives in harsh coastal environments. Unlike structures derived from laboratory grown specimens, our analysis captures PSI directly from wild-grown plant leaves, offering an in-situ snapshot of plant adaptation. The core architecture comprising twelve protein subunits and four Lhca antenna proteins coordinating 157 chlorophylls, 36 carotenoids and 24 lipids is highly conserved relative to model angiosperms yet exhibits subtle shifts in antenna core coupling and pigment organization. Red-shifted chlorophylls in Lhca3, conserved among far red adapted plants, suggest an ancestral photoprotective mechanism retained in high light, high-salinity environments. Lipid mediated stabilization at interfaces likely counteracts salinity-induced membrane fluidity changes enhancing resilience under fluctuating light and salt stress. Comparative analysis with PSI structures from other vascular plants highlights both evolutionary conservation and distinct ecological tuning of LHCI organization and cofactor binding in mangroves. These insights reveal molecular strategies enabling *A. marina* to maintain efficient photosynthesis under extreme conditions, offering strategies for engineering photosystems in crops.

## Introduction

Since the inception of photosynthesis, phototrophic organisms have successfully colonized a wide array of environmental niches, ranging from the depths of oceans to dry deserts (Garcia-Pichel et al., 2001; Gattuso et al., 1998). Their remarkable adaptability to diverse conditions is attributed to their ability to convert light of varying quality and intensity into biochemical energy (Annemarie et al., 2024). Photosynthesis harnesses light energy to synthesize organic compounds from inorganic precursors, specifically converting carbon dioxide and water into carbohydrates, which are the building blocks of biomass while releasing molecular oxygen in the atmosphere and an equivalent reduction in carbon dioxide levels (Blankenship, 2002; Calvin & Benson, 1948). This process is predominantly carried out by oxygenic photosynthetic organisms such as plants, algae, and cyanobacteria. The primary reactions of photosynthesis occur within the thylakoid membranes of cyanobacterial cells and the chloroplasts of eukaryotic organisms, a specialized organelle arising from endosymbiotic integration of cyanobacterial ancestors (Zak et al., 2001). Structurally, chloroplasts are surrounded by a double membrane envelope, a feature consistent across plant and algal species (Block et al., 2007). Within these chloroplasts, the thylakoid membrane contains the multisubunit pigment protein complexes responsible for the photosynthetic electron transport chain (Dekker & Boekema, 2005). Thylakoids are arranged in stacked formations called grana, interconnected by stromal lamellae, creating an extensive network for light-dependent reactions (Nelson & Junge, 2015). Photosystem II (PSII) is enriched in grana stacks, whereas PSI predominantly localizes to stroma-exposed regions, where it can interact with soluble electron carriers. This structural organization ensures efficient light energy capture and conversion into biochemical energy (Nelson & Ben-Shem, 2005; Pribil et al., 2014).

PSII initiates the light dependent reactions of photosynthesis by capturing photon energy to catalyze the oxidation of water molecules (Barber, 2016). This process results in the release of molecular oxygen and the extraction of electrons, which are subsequently transferred to the plastoquinone (PQ) pool (Shen, 2015). Concurrently, protons generated from water splitting, along with additional protons translocated by the cytochrome b₆f complex, accumulate within the thylakoid lumen (Renger & Renger, 2008). The resulting proton gradient across the thylakoid membrane establishes a proton motive force that drives ATP synthesis via the membrane-bound ATP synthase complex, coupling the conversion of ADP and inorganic phosphate into ATP with the dissipation of the electrochemical gradient (Maloney, 1982). Conversely, PSI absorbs photons of longer wavelengths to elevate electrons from plastocyanin to ferredoxin (Nelson & Junge, 2015). These high-energy electrons are then shuttled through a chain of redox-active carriers, culminating in the reduction of NADP⁺ to NADPH (Johnson, 2011). NADPH, along with ATP, serves as a key reductant and energy currency for the subsequent carbon fixation reactions in the Calvin-Benson cycle (Blankenship, 2014; Barber, 2009).

The core components of photosystems are composed of numerous protein subunits and a diverse array of cofactors, including chlorophylls, carotenoids, quinones, and lipids. Every photon absorbed by PSI drives the transfer of one electron, giving PSI an efficiency equal to ≈ 100 % (Nelson & Yocum, 2006).

In plants, PSI is a large macromolecular complex with a molecular weight of approximately 700 kDa. The structures of PSI from various organisms have been determined at high resolution by X-ray crystallography, providing insights into their electron transfer mechanism and function (Amunts et al., 2007; Ben-Shem et al., 2003; Mazor et al., 2017; Qin et al., 2015). While the basic building blocks of photosynthesis are conserved, modular pigment protein complexes have developed in different organisms to maximize the sunlight driven energy conversion. Therefore, considerable variation exists in the architecture of thylakoids and their peripheral antenna complexes (Shen, 2022). The plant PSI structure comprises core and light-harvesting (LHCI) complexes, which together form the PSI-LHCI supercomplex (Ben-Shem et al., 2003). The PSI core complex is typically divided into two large subunits - PsaA and PsaB. Both contain special chlorophylls forming the primary electron donor P700 and primary acceptors, phylloquinones, and an iron-sulfur [4Fe-4S] cluster involved in the electron transport chain (Jensen et al., 2007). PsaA/B also binds most of the chlorophylls and cofactors. The cytoplasmic PsaC subunit is stabilized by its neighbour PsaD and PsaE, anchors additional two [4Fe-4S] clusters reducing ferredoxin, while the remaining small transmembrane subunits play an important role in the function and organization of PSI. Previous studies indicated that green algal and diatome PSI-LHCI is more varied than that of plants (Bai et al., 2021). Plant PSI-LHCI typically comprises four light-harvesting proteins, Lhca1-4, and their positions generally correspond to a conserved configuration (Croce & Van Amerongen, 2013). These Lhca proteins bind numerous chlorophyll molecules and carotenoids, significantly contributing to the light absorption capabilities of the PSI (Caspy & Nelson, 2018).

Global climate change represents one of the most urgent challenges of the 21st century and a significant threat to the stability and resilience of Earth’s ecosystems (Malhi et al., 2020). Photosynthesis plays a central role in regulating Earth’s climate (Tkemaladze & Makhashvili, 2016). However, the accelerating pace of global warming is disrupting photosynthetic functions across different ecosystems from forests to phytoplankton blooms (Hallegraeff, 2010). To fully understand and predict the adaptive responses of photosynthesis under these changing conditions, it is essential to study this process not only within controlled laboratory environments, but also in wild plants. In the context of climate change resilience, mangroves are essential to protect coastlines and cities built at the sea, as they form a natural barrier against inundation (Spalding, 2010). They demonstrate remarkable adaptability, where they face a combination of stressors including high salinity, nutrient deficiency, waterlogging, and intense light exposure. The majority of plant species could not survive such harsh conditions (Ball & Critchley, 1982). Mangrove ecosystems occupy a mere 0.36 % of the world’s forested area, yet account for nearly 10–15 % of coastal marine carbon stocks. They store roughly 6–12 petagrams (Pg; 1 Pg = 10¹⁵ g) of carbon in their biomass and have sediment rates of sequestration that are four to ten times those of terrestrial rainforests on an area basis (Kauffman et al., 2020; Wang et al., 2025). These halophytic woodlands fringe tropical and subtropical shorelines, where tidal inundation, hypersalinity (up to ∼600 mM NaCl), waterlogging, and supra-optimal irradiance (> 2 000 µmol photons m⁻² s⁻¹) impose a suite of concurrent stresses on photosynthetic machinery (Chen et al., 2022). Field measurements reveal that mature mangrove trees of the species *Avicennia* and *Rhizophora* can sustain midday gross photosynthetic rates of 18–22 µmol CO₂ m⁻² s⁻¹ and maintain photochemical quantum yields (Fv/Fm ≈ 0.78–0.82) despite leaf-surface temperatures exceeding 45 °C and xylem Na⁺ concentrations above 500 mM (Cheeseman & Lovelock, 2004). These performance metrics rival and in situ often surpass those of C₄ crops under controlled-environment conditions (Chatting et al., 2022; Mcleod et al., 2011), suggesting sophisticated physiological mechanisms for sustaining photosynthetic activity under potentially photoinhibitory conditions.

*Avicennia marin*a, known regionally as Qurm in the Arabian Gulf, the most abundant mangrove species (Haseeba et al., 2025) thrives under extreme salinity (up to 100 PSU), intense solar radiation, and arid conditions through specialized adaptations including salt excreting glands, osmotic regulation, and pneumatophores (Adame et al., 2021). Its success in such a challenging habitat makes *A. marina* a valuable model for studying plant resilience in multi stress environments.

In this study, we present the structural analysis of the PSI–LHCI supercomplex from *A. marina*. The samples were collected from Jubail Ali Mangrove Park in it’s natural environments (UAE: 25.0°N, 55.1°E). Photosynthetic membrane complexes were isolated and their structure was determined using single-particle cryo-electron microscopy (cryo-EM), resulting in a reconstruction at an overall resolution of 2.05 Å. The PSI core exhibits a conserved structural organization similar to that found in higher plants. The LHCI antenna also shows a conserved arrangement compared to other plants, with four Lhca1-4 proteins bound. We identified several lipid molecules bridge the antenna-core interface, stabilising the complex and potentially tuning excitation equilibration.

## Results

To gain structural insight into the light-harvesting apparatus of mangrove plants under natural environmental stress, we determined a high-resolution cryo-EM structure of the PSI–LHCI supercomplex from *A. marina*, a halotolerant mangrove species. A unique aspect of this study is the origin of the sample. Unlike most structural investigations that rely on controlled laboratory growing conditions, we extracted intact thylakoid membranes directly from field-collected *A. marina* leaves near Abu Dhabi in the Arabian Gulf (**Extended Fig. 1**).

### Overall architecture and subunit composition

To determine the structure of the *A. marina* PSI–LHCI supercomplex, we employed single-particle cryo-electron microscopy on native thylakoid membrane extracts, ultimately resolving the complex at 2.1 Å resolution. Extensive classification and refinement across CryoSPARC and RELION platforms yielded a final dataset of 157,448 high-quality particles (Extended Data Fig. 2–3; see Methods for details). The overall architecture of the *A. marina* PSI–LHCI supercomplex conforms to the canonical design observed in higher plants such as *Pisum sativum*, *Arabidopsis thaliana*, *Zea mays*, and *Spinacia oleracea*, underscoring the structural conservation across angiosperms despite the extreme environmental pressures encountered by halotolerant species. The PSI core in our structure comprises 12 subunits, spanning PsaA through PsaL, and four Lhca antenna proteins (Lhca1–Lhca4), alongside 156 chlorophylls, 36 carotenoids, and 25 lipid molecules (**Fig. 1a, b**).

**Figure 1.**
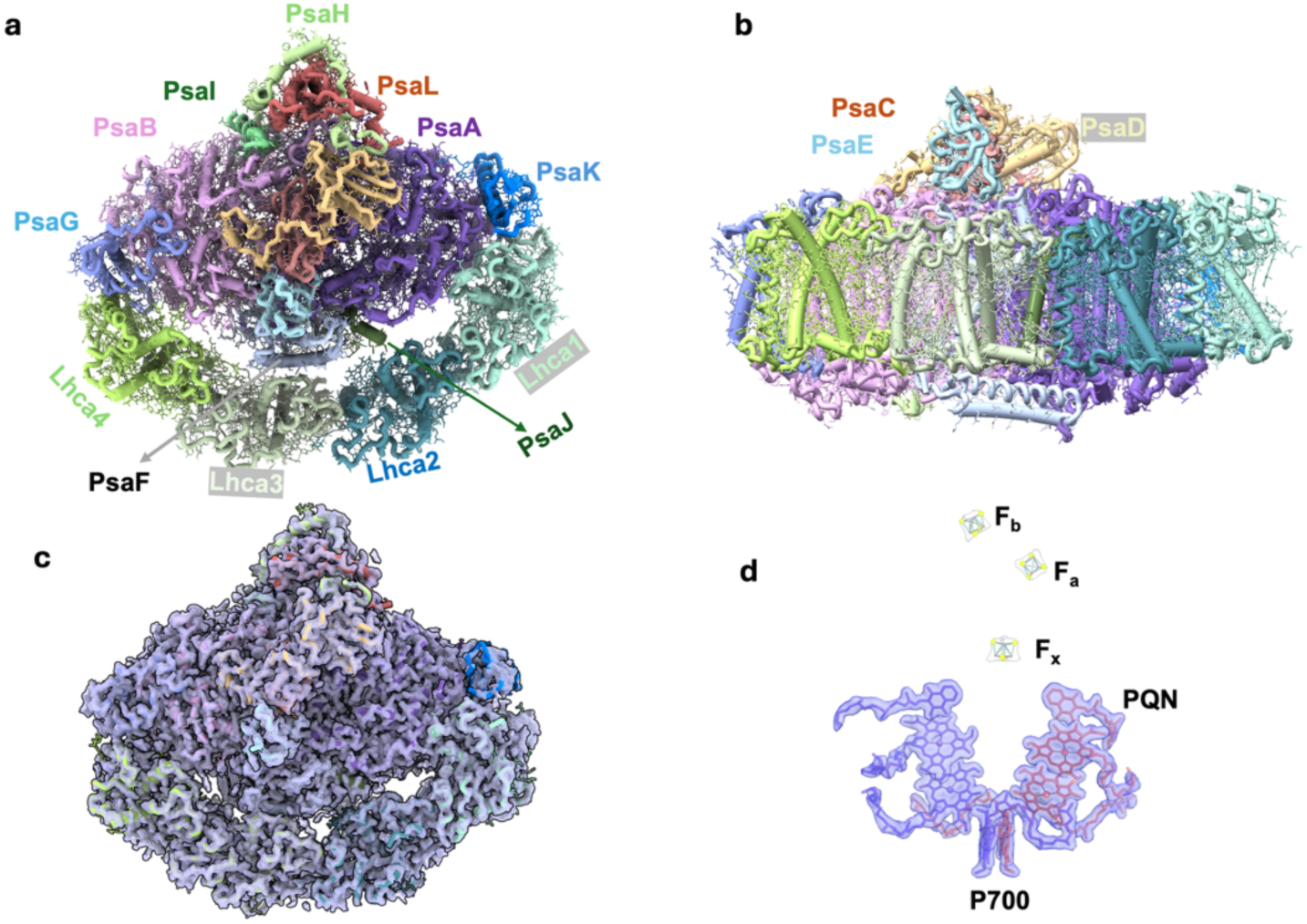
Overall architecture and subunit composition of PSI-LHCI supercomplex. **a)** Top view of the architecture of *A. marina* PSI–LHCI supercomplex, showing structural conservation with other angiosperms such as *Pisum sativum*, *Arabidopsis thaliana*, *Zea mays*, and *Spinacia oleracea*. **b)** Side view of *A. marina* PSI–LHCI supercomplex. **c)** Overlay of structural model with electron density at approximately 2.05 Å. **d)** Structure of electron transfer chain cofactor with density of P700, PQN and iron-sulfur clusters Fx, F_A_ and F_B_.

Notably, we did not observe the density corresponding to the PsaO subunits likely because this subunits only binds to PSI-LHCI during state transition (state 2) together with a phosphorylated LHCII trimer (Wu et al.,2023), nor the density associated with PsaN likely lost during purification, similar to other PSI LHCI structures (Mazor et al., 2015) (**Fig. 1c**).

The PSI core exhibits a conserved arrangement of nine transmembrane subunits (PsaA–PsaL) and three stromal extrinsic subunits (PsaC, PsaD, PsaE). The heterodimeric reaction center formed by PsaA and PsaB coordinates the electron transport cofactors, including the special pair chlorophylls (P700), the primary electron acceptors (A₀), phylloquinones (A₁), and the iron-sulfur cluster (FX). The stromal ridge formed by PsaC, PsaD, and PsaE houses the terminal iron-sulfur clusters FA and FB, (**Fig. 1d**). The PSI core contains 96 chlorophyll *a* molecules that are densely interspersed among the nine to ten transmembrane α-helices contributed by both major and minor subunits.

Structural alignment with PSI–LHCI complexes from *Pisum sativum* (PDB: 7DKZ), *Hordeum vulgare* (7EWK), *Zea mays* (PDB:5ZJI), *Spinacia oleracea (*PDB:9GRX*), Avena sativa (*PDB:8BCV), and *Fittonia albivenis* (PDB:8WGH) reveals minimal deviations (Cα RMSD <1.0 Å), affirming evolutionary conservation of the core fold (**Extended Fig. 4a**) (Introini et al., 2025; Li et al., 2024; Naschberger et al., 2024; Pan et al., 2018; Shen et al., 2022; Wang et al., 2021). PsaF, PsaI, and PsaJ each consist of a single transmembrane helix, PsaG and PsaK contain two transmembrane helices, and PsaL comprises three transmembrane helices. PsaG and PsaK are structurally similar, particularly in their transmembrane helices. In our structure, the PsaG subunit is well resolved. However, around ten residues of the subunit PsaK first transmembrane helix remain poorly resolved loop, consistent with its known flexibility across species. This conformational variability could result in the adaptation to light, temperature, and salinity fluctuations during plant growth reflecting PsaK’s dynamic role in natural plant growth conditions (Mazor et al., 2017). PsaF protrudes on the lumenal side near the PsaA/PsaB interface, contributing to the docking site for plastocyanin, while PsaD and PsaE form a protruding ridge on the stromal side that binds ferredoxin/flavodoxin for electron transfer (Caspy et al., 2020; Finazzi et al., 2005).

### LHCI organization and excitation energy transfer

The four LHCI antenna proteins (Lhca1–Lhca4) in *A. marina* PSI–LHCI form a crescent-shaped belt (**Fig. 2a, b**) that surrounds one side of the core complex (Li et al., 2024; Novoderezhkin & Croce, 2023; Qin et al., 2015; Zhang et al., 2023). This arrangement is analogous to the canonical configuration observed in other angiosperms, including *A. thaliana, P. sativum*, and *S. oleracea*. Unambiguous assignment of Lhca1–Lhca4 in the *A. marina* complex was achieved based on the cryo-EM density and guided by ModelAngelo-assisted model building. The four antenna subunits were confidently assigned (despite limited sequence information for *A. marina*) by structure and sequence alignments to known LHCI proteins. Each Lhca is a three-helix membrane protein, binding chlorophyll a/b and carotenoids, and they attach in series along the PsaF/PsaG/PsaK side of the core (**Fig. 2a**).

**Figure 2.**
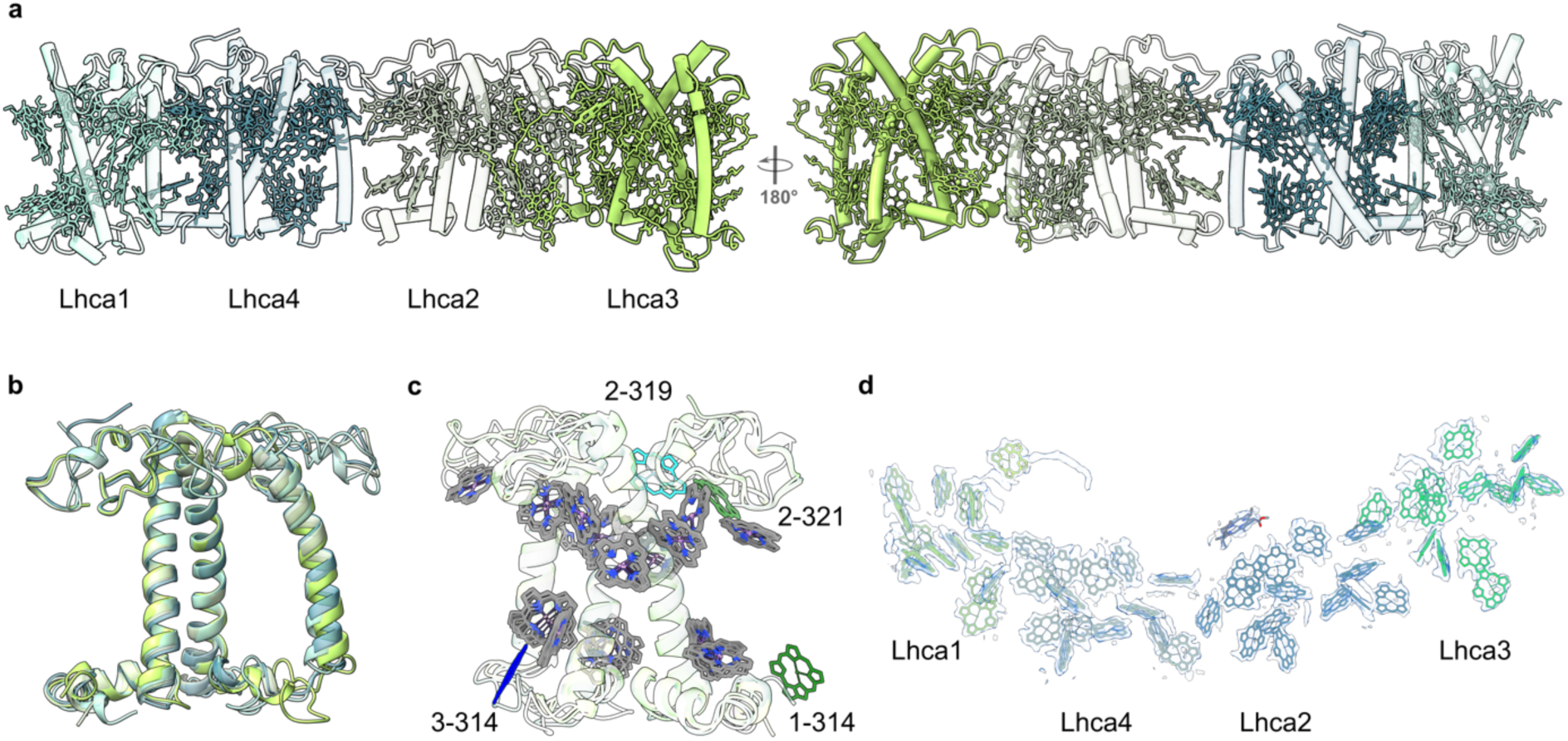
LHCI Organization and Excitation Energy Transfer. **a)** Structural representation of Lhca1– Lhca4 subunits of *A. marina* PSI-LHCI, visualized parallel to the membrane plane from both stromal (inner) and lumenal (outer) perspectives. The apoprotein backbones and associated pigments are depicted in stick representation, highlighting the spatial arrangement of chlorophyll and carotenoid cofactors within each subunit. **b)** Superimposition of the four Lhca apoprotein structures reveals conserved and divergent elements within the Lhca. **c)** Overlay of chlorophyll cofactors from all four Lhca subunits, with unique chlorophyll positions distinguished by color (Lhca1: green, Lhca2: cyan, Lhca3: blue), delineates both shared and subunit-specific pigment-binding sites that may underpin functional specialization in light-harvesting and energy transfer. **d)** Density map for chlorophyll molecules.

The order of Lhca around the PSI core is Lhca1–Lhca4–Lhca2–Lhca3, consistent with the canonical arrangement in higher plants (Castelletti et al., 2003; Chen & Blankenship, 2011). The inter-subunit contacts, particularly the Lhca1–PsaG and Lhca3–PsaK interfaces, involve conserved amino acid interactions that appear to be evolutionarily preserved across diverse plant lineages. These conserved interfaces ensure stable association between the core and antenna domains, which is essential for efficient excitation energy transfer. Indeed, the pathways for excitation energy transfer appear preserved, as evidenced by the conserved spatial arrangement of bridging chlorophylls between the LHCI belt and PSI core (Introini et al., 2025; Li et al., 2024; Naschberger et al., 2024; Pan et al., 2018; Qin et al., 2015; Wang et al., 2021; Zhang et al., 2023). Superposition of all four Lhca subunits showed their high overall structural similarity (**Fig. 2c**). However, notable deviations were observed in the first and third transmembrane loop regions. Chlorophyll-binding sites are largely conserved in position upon alignment, though chlorophylls 314 and 321 exhibit displacement relative to their counterparts (**Fig. 2d**). In our structure, their slightly altered orientations and inter-pigment distances may reflect functional tuning to fluctuating environmental stress in natural growth conditions.

Despite strong structural conservation, several species-specific features distinguish the *A. marina* PSI–LHCI complex from models such as *A. thaliana*, *Z. mays* or *S. oleracea* A key feature is the Mg–Mg distances between the P700 special pair and LHCI chlorophylls (e.g., Lhca1: 63.9 Å, Lhca2: 55.3 Å, Lhca3: 75.6 Å, Lhca4: 58.1 Å), which align closely with values reported for shade adapted species, suggesting preserved excitation energy transfer pathways. Notably, Lhca3 retains a red-shifted chlorophyll pair (Chl a303/a309), stabilized by Asn-Mg coordination as like in most angiosperm, but with a protein comprising conserved aromatic residues like shade adapted species (Li et al., 2024). These features suggest a shared mechanism for low-energy chlorophyll stabilization and flexible LHCI–core coupling under stress across angiosperms. If the geometry of the PSI core remains largely conserved, exhibiting no significant structural rearrangements, minor lateral displacements are observed within the LHCI belt (RMSD ∼0.3–0.5 Å), indicative of flexible antenna–core coupling that may facilitate adaptation under environmental stress conditions (**Extended Fig. 4a, b**).

Our PSI–LHCI reveals a total of 241 prosthetic groups. Of these, 156 were modeled as chlorophyll pigments, including 12 chlorophyll b molecules localized within the light-harvesting complexes. Accessory pigments comprised 36 carotenoids, specifically 25 β-carotenes, 6 luteins, and 5 xanthophylls. In addition, 24 lipid molecules were identified, including 21 monogalactolipids (12 LMG and 4 LHG), 5 digalactolipids (DGD), and 4 phospholipid derivatives, all of which contribute to the structural stabilization of the complex. Pigment binding sites are highly conserved when compared with previously resolved plant PSI structures (**Fig. 3a**). Structural superposition of *A. marina* PSI–LHCI (green) with other plant PSI models (yellow, PDB 8WGH; magenta, PDB: 7EWK; orange, PDB: 5ZJI) (**Fig. 3b, c**) demonstrates strong conservation of chlorophyll and carotenoid positions across species. Notably, our model reveals a unique chlorophyll located at the interface between Lhca2 and PsaF (**Fig. 3c**-green), which may represent an additional excitation energy transfer pathway bridging the antenna and core complexes.

**Figure 3.**
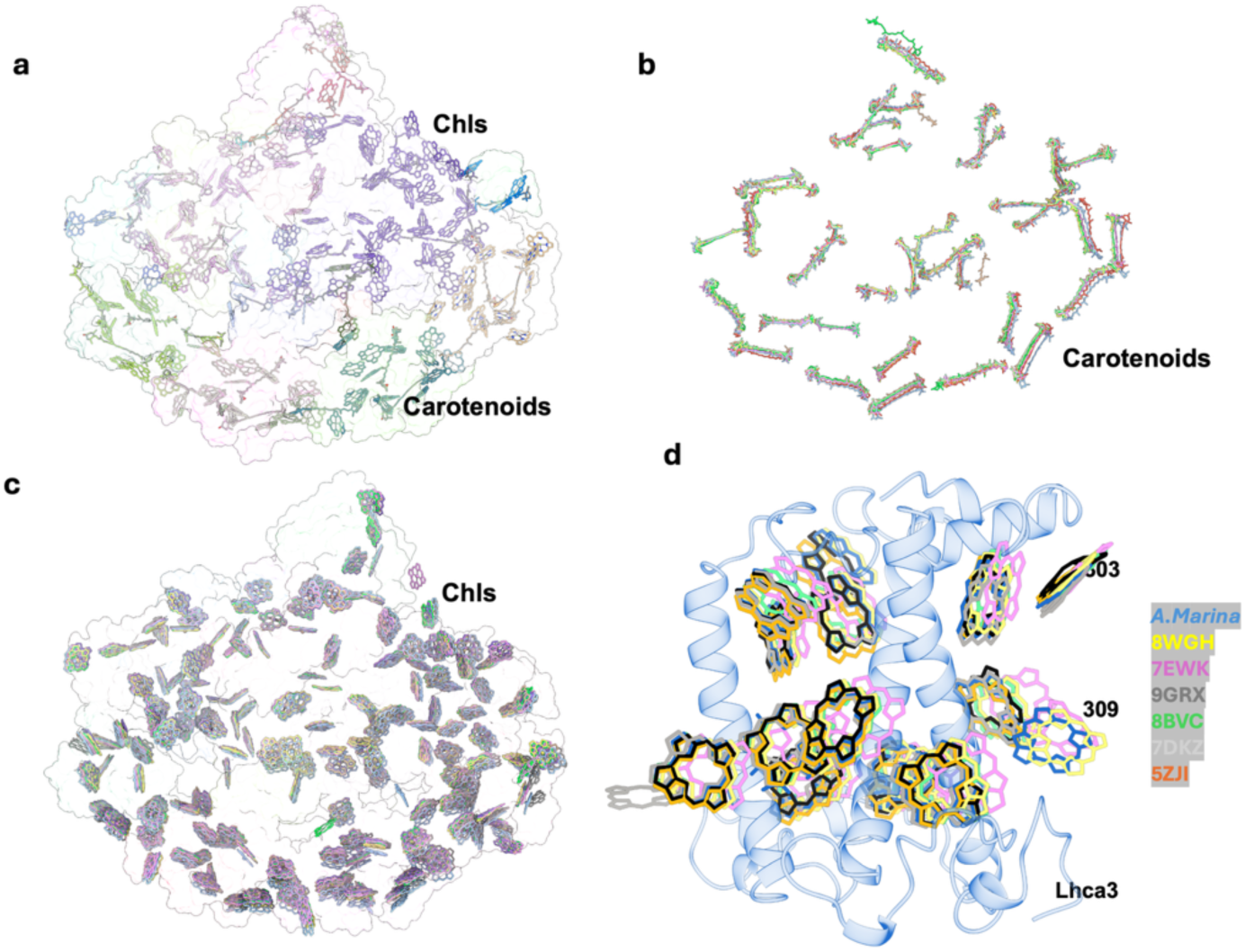
Distribution and Comparative Analysis of Carotenoids and Red Chlorophylls in *A.marina* PSI–LHCI. **a)** Spatial arrangement of chlorophylls and carotenoids in the *A. marina* PSI–LHCI complex, with pigment positions colored according to their respective subunits. For clarity, phytol chains have been omitted from the chlorophyll models. **b)** Superposition of carotenoid positions in *A. marina* with those from other plant PSI structures reveals a high degree of spatial conservation, underscoring the evolutionary stability of carotenoid binding sites. **c)** Comparative mapping of chlorophyll positions across plant PSI structures. **d)** Detailed comparison of Lhca3-associated chlorophyll positions among various plant PSI– LHCI complexes: *A. marina* (blue), 8WGH (*F. albivensis* in yellow), 7EWK (*H. vulgare* in magenta), 9GRX (*S. oleracea* in dark), 8BCV (*A. sativa* in green), 5ZJI (*Z. mays* in orange), and 7DKZ (*P. sativum* in light grey).

While *F. albivenis* employs extended Lhca3 loop regions to enhance red-shifted absorption, *A. marina* retains canonical loop conformations suggesting similarity with e.g *A. thalina* and that *A. marina* far-red chlorophylls serve for far-red light absorption under full sun light rather than other niche-specific adaptation like e.g shade adaptation (Li et al., 2024).

Because we shown that *A. marina* display feature in the LHCI that are similar to the shaded plant *F. albivensis* and To further elucidate the structural adaptation of light harvesting, we compared of the Lhca3 subunit with those from previously reported plant PSI–LHCI structures(PDB IDs: 5ZJI, 7DKZ, 7WFD, 8BCV, 8WGH, and 9GRX) (**Fig. 3d** & **Extended Fig. 6b**). Our model reveals that the positions of red-shifted chlorophylls within *A. marina* Lhca3 are remarkably consistent with those identified in the far-red light-acclimated Ficus albivenis PSI–LHCI structure.

Superposition analyses show that these red chlorophylls occupy analogous binding pockets and exhibit comparable coordination environments, indicating a conserved structural motif among species adapted to challenging light conditions. Furthermore, we observed that the configuration of red chlorophylls in our *A. marina* model aligns closely with those found in Lhca5 in the the NDH-associated cyclic electron transport PSI complex of *A. thaliana* (Su et al., 2022).

This structural congruence extends to both the spatial orientation and the protein environment of the chlorophyll ligands, suggesting that the stabilization of red-shifted pigments in *A. marina* is a feature shared among plant lineages experiencing shaded and far-red enriched light like *F. albivensis.* (**Fig. 3d**).

Carotenoids and lipid variations may reflect habitat-specific photoprotection and stabilization, with *A. marina* maintaining 36 carotenoids compared to 35–38 in other plants, suggesting sufficient built-in mechanisms without additional pigment recruitment in PSI-LHCI complex. In our structure we model 25 Lipids were assigned according to the density map (**Fig. 4a, b**) also assigned several detergent molecules. Lipids interactions, particularly at core–antenna interfaces, likely counteract membrane fluidity changes under salinity, a feature less pronounced in laboratory-grown *A. thaliana*. At the PSI–LHCI interface, three lipid molecules were resolved with sufficient clarity to permit modeling in the unique position comparing with other plant PSI structure (**Fig. 4c**). These lipids were predominantly located at the PsaG-Lhca1, PsaF–Lhca2 and PsaK–Lhca3 junctions. Their acyl chains inserted into hydrophobic clefts between transmembrane domains, while polar headgroups formed stabilizing interactions with adjacent protein residues. For example, one LHG molecule bridges between the lumenal face of PsaF and Lhca1, anchoring the antenna subunit via hydrogen bonding which is consistent with previous reports (Croce & Van Amerongen, 2014; Davison et al., 2002; Dekker & Boekema, 2005; Havaux, 1998; Mazor et al., 2017). These lipid interactions are consistent with previous findings in *P. sativum* and *A. thaliana* PSI–LHCI complexes, where lipids were shown to facilitate supercomplex assembly and maintain subunit integrity. The conservation of these lipid-binding sites in *A. marina*, despite its saline growth environment, suggests that membrane lipid–protein interactions play a critical role in PSI structural stability across diverse taxa. This lipid-mediated stabilization is conserved across plant species, suggesting its fundamental importance. In the saline mangrove environment, where high salinity could disrupt membrane fluidity, these interactions may be particularly critical for maintaining the structural integrity of the photosynthetic apparatus under stress. A recent structural analysis of the extremophilic green alga *C. ohadii* revealed that carotenoids and lipids play critical roles in photoprotection and in stabilizing the photosynthetic complex under harsh environmental conditions. These components appear to be integral to the organism’s ability to maintain photosystem integrity in extreme habitats (Caspy et al., 2021; Nelson, 2024).

**Figure 4.**
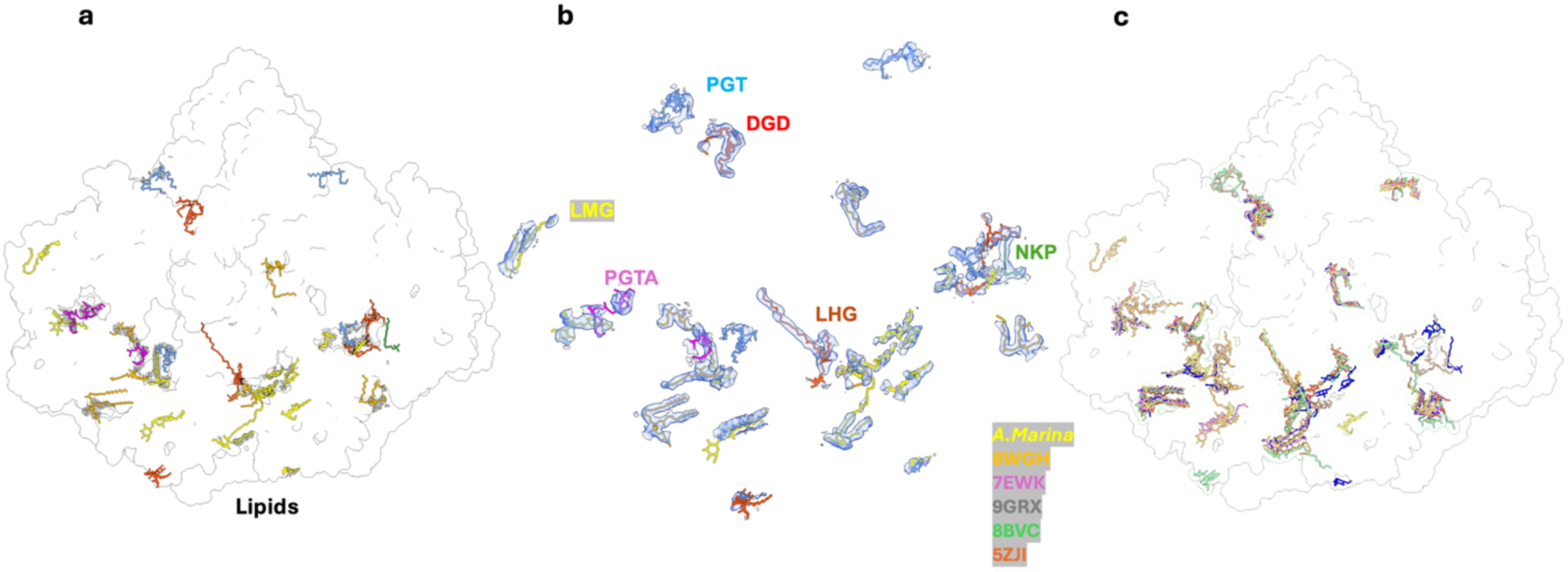
Distribution and Conservation of Lipid Molecules in A. marina PSI–LHCI. **a)** Spatial distribution of lipid molecules within the *A. marina* PSI–LHCI complex. Lipids are shown associated with specific subunits, with different lipid types distinguished by color. **b)** Cryo-EM density map highlighting the positions and assignments of lipid molecules within the PSI– LHCI-supercomplex. **c)** Structural superposition of lipid molecules from A. marina (yellow) with those from other plant PSI–LHCI structures: (orange, PDB: 8WGH), (magenta, PDB: 7EWK), (dark, PDB: 9GRX), and (green, PDB: 8BCV). The high degree of spatial conservation underscores the functional significance of lipid–protein interactions in PSI assembly and stability.

## Discussion

The determination high resolution of the 2.1 Å cryo-EM structure of the PSI–LHCI supercomplex from *A. marina*, a halotolerant mangrove species, provides key insights into the structural resilience and evolutionary conservation of photosystem I under naturally fluctuating and physiologically stressful conditions. The final map reveals a comprehensive structural model of the PSI–LHCI complex, with sufficient clarity to resolve individual amino acid side chains and bound pigments. This study represents one of the few instances in which a plant photosystem has been resolved directly from field-collected material, offering a uniquely authentic view of how photosynthetic machinery operates in situ, without the potential introduced by control growth environments. The architecture of the complex confirms a high degree of structural conservation across vascular plants, while also revealing specific adaptations likely linked to the unique intertidal habitat.

### Ecological and Evolutionary Implications

The structural conservation of the PSI–LHCI supercomplex across angiosperms, despite their divergent habitats, reflects the evolutionary optimization of this photosynthetic design. However, the subtle variations in *A. marina*, such as the absence of PsaN/PsaO, minor LHCI shifts, and conserved red-shifted chlorophylls, indicate fine-tuning to its specific environmental challenges. Furthermore, the subtle positional shifts in the LHCI belt and the slightly longer inter-chlorophyll distances in *A. marina* compared to model plants are not random variations. These minor structural deviations could influence the kinetics and efficiency of excitation energy transfer (EET) from the antenna to the reaction center. Slightly longer distances might lead to marginally slower EET rates, which could be an adaptive strategy to enhance photoprotection under high-light stress by allowing more time for non-photochemical quenching, balancing efficient light harvesting with robust photodamage prevention in exposed intertidal zones. The observed flexibility in LHCI-core coupling and the positional shifts in the LHCI belt suggest that the PSI–LHCI supercomplex is not a static entity but possesses intrinsic dynamic properties. This flexibility might allow dynamic adjustments to optimize energy transfer pathways under fluctuating light conditions, or to accommodate interactions with other regulatory proteins (e.g., during state transitions where LHCII might transiently associate).

Taken together, our comparative analysis highlights a possible convergence in pigment architecture between mangrove and far-red light-adapted species. This is surprising given the natural habitat of *A. marina* (high light/temperature and salinity) compared to *F. albivenis* (shaded and far red enriched light). One hypothesis is that the energetic demands of *A. marina*, like salt stress resistance, may have driven an increase in far-red light harvesting, leading to evolutionary convergence with plants adapted to shaded, far-red-enriched environments. The fact that *A. marina* exhibits unique features in a wild-collected state indicates that these are adjustments as part of its natural physiological response to environmental variability (Oku et al., 2003). These findings emphasize the value of field-derived structural data in revealing ecologically relevant adaptations that might be missed in laboratory-controlled settings. By providing a high-resolution snapshot of photosynthesis in a stress-tolerant plant, this study bridges structural biology and ecological physiology.

The *A. marina* PSI–LHCI structure reveals a photosynthetic apparatus that may be conserved and subtly adapted to extreme conditions. The minor structural differences likely enhance photoprotection and stability, enabling the plant to thrive in its harsh intertidal habitat. Key innovations such as lipid-mediated stabilization, and flexible antenna-core coupling highlight mechanisms that reconcile efficient energy transfer with photoprotection. These insights not only expand our understanding of photosystem evolution but also inform strategies for engineering stress-tolerant crops in saline or marginal environments.

In conclusion, the *A. marina* PSI–LHCI provides new insights for how photosynthetic complexes can be adapted to extreme environments. Its structural innovations in pigment arrangement and lipid anchoring illuminate the molecular basis of enhanced photoprotection. The preservation of key photochemical elements, alongside minor adaptive shifts in pigment organization and protein– lipid interactions, exemplifies a sophisticated balance between evolutionary conservation and environmental modulation that ensures optimal photosynthetic performance in an extreme coastal habitat.

## Materials and methods

### Plant Material

Approximately 200 grams of fresh *Avicennia marina* leaves were harvested during morning hours from Jubail Mangrove Park, Abu Dhabi (24°32′48.07″N, 54°29′14.78″E). Leaves were washed with distilled water to eliminate surface salt secretions. Approximately 100–150 grams of this fresh leaf tissue were then prepared for homogenization. Isolation of thylakoid membranes and PSI from mangrove leaves was performed according to previously established protocols (Ball et al., 1984; Caspy et al., 2020; Charras et al., 2024), with modifications. Mangrove leaf tissue was gently pressed into a chilled blender and covered with a modified ice-cold homogenization buffer contain of 0.33 M sucrose, 25 mM Tris-NaOH (pH 7.5), 15 mM NaCl, 5 mM MgCl₂, 1 mM PMSF, 1 mM benzamidine, and 5 mM aminocaproic acid and 1 mM EDTA supplemented with 0.15% (w/v) bovine serum albumin (BSA), 4 mM sodium ascorbate, and 7 mM cysteine to prevent oxidative degradation. Batches of 30–35 g of leaves were processed in a chilled blender using a sequence of 10 s on, 10 s off, and 10 s on to limit heat buildup. The homogenate was filtered through four layers of 20 μm nylon mesh to remove debris. Intact chloroplasts were then isolated via a 40–80% Percoll step gradient, briefly, 80% Percoll was carefully layered beneath 40% Percoll buffer, maintaining a distinct interface. The homogenate was overlaid on the gradient and centrifuged, with intact chloroplasts collecting at the 40–80% boundary.

### Isolation and Purification of PSI

All extraction and purification steps were conducted at 4°C under dim illumination to minimize chlorophyll photodegradation. Chloroplast pellets were resuspended in 1 L of hypotonic buffer, using gentle pipetting or swirling, followed by homogenization with a glass tissue grinder. Thylakoid membranes were isolated by centrifugation of the homogenate at 1,000 × g for 9 min. The resulting pellet was resuspended in STN buffer (0.4 M sucrose, 50 mM Tricine, pH 7.8, 10 mM NaCl) to yield a final chlorophyll concentration of 1.5 mg/mL, as determined spectrophotometrically following dilution in 80% (v/v) acetone. For PSI isolation, the thylakoid suspension was solubilized with n-dodecyl-β-D-maltoside (α-DDM) at a detergent-to-chlorophyll mass ratio of 15:1 and incubated on ice for 30 min. The solubilized material was clarified by ultracentrifugation at 120,000 × g for 30 min. The supernatant was loaded onto a Toyopearl DEAE ion-exchange column pre-equilibrated in buffer, and PSI was eluted using a linear NaCl gradient (0–250 mM). PSI-containing fractions were identified spectroscopically, pooled, and concentrated. Subsequently, the pooled PSI fractions were layered onto a 10–30% (w/v) sucrose step gradient in STN buffer and subjected to ultracentrifugation at 100,000 × g for 20 h using an SW-40 rotor. The distinct dark green PSI band was collected, concentrated using a 100 kDa molecular weight cutoff concentrator, and subjected to buffer exchange to remove sucrose. For final purification, the sample was applied to a continuous sucrose gradient (15–50%, w/v) prepared in buffer containing 20 mM Tris-Tricine (pH 8.0) and 0.05% α-DDM, followed by centrifugation in an SW-60 rotor at 55,000 rpm for 15 h. The purified PSI complex appeared as a discrete dark band, which was harvested for subsequent analysis. The migration position of the PSI band in the sucrose gradient is indicated in Extended Fig. 1j (asterisk), corresponding to one of twelve gradient fractions. The second fraction, containing the enriched PSI complex, was collected and used for all further analyses.

### Gel Electrophoresis

To analyze the polypeptide composition of isolated PSI complexes, samples were treated with sodium dodecyl sulfate (SDS) sample buffer containing 2% (w/v) SDS, 60 mM dithiothreitol, and 60 mM Tris-HCl (pH 8.5). The mixture was heated at 40 °C for 10 min to ensure complete denaturation. Subsequently, the samples were subjected to SDS-polyacrylamide gel electrophoresis (SDS-PAGE) as described by Ikeuchi and Inoue, using a 16% polyacrylamide gel (acrylamide:bisacrylamide ratio of 37.5 : 1 [w/w]) supplemented with 7.5 M urea to resolve chlorophyll-binding proteins. Samples corresponding to 2.5 µg of chlorophyll (Chl) were loaded per lane to ensure consistent protein quantification.

### Absorption Spectroscopy

Absorption spectra of mangrove leaves and isolated PSI samples were recorded at room temperature using a UV-2700i Plus UV-Vis Spectrophotometer (Shimadzu, Japan) across the wavelength range of 350–750 nm. Isolated samples were diluted in a buffer containing 10 mM Tricine-NaOH (pH 7.5) and 0.03% (w/v) α-DDM to achieve a chlorophyll concentration with a maximal absorption peak of approximately 0.8. Baseline correction was performed by setting absorption at 750 nm set to zero.

### Cryo-EM image processing

12,252 micrographs were imported into CryoSparc and motion-corrected (Punjani et al., 2017). The blob-picker picked 2,292,198 particles at 500 pixels box size and binned to 100 pixels. We performed two rounds of 2D classification to filter particles that were not PSI. Selected 2D classes were used for ab initio, re-extraction at 500 pixels, non-uniform refinement and heterogenous refinement. 3D classification into five classes yielded classes of different quality. We picked the highest quality class of 611,477 particles, performed another non-uniform refinement, generating a map of **2.94** Å and exported the particles for further processing in Relion.

611407 particles were imported into Relion5 (Burt et al., 2024). First, a 3D refinement was performed and the resulting map used to create a volume in ChimeraX, which was subsequently made into a solvent mask in Relion. This mask was used for all subsequent refinements. A second 3D refinement was followed up by post-processing, CTF refinement (beamtilt, anisotropic and parameter fitting) and polishing, yielding a map of **2.41** Å (**Extended Fig. 2**). We then performed a third round of 3D refinement, post-processing, CTF refinement (CTF with 4^th^ order aberration, anisotropic and parameter fitting), leading to a map of **2.38** Å. This was followed up by the 4^th^ 3D refinement (**2.37** Å), post-processing (**2.38 Å**), polishing, the 5^th^ 3D refinement (**2.34 Å**) and post-processing (**2.34 Å**). As the resolution improvement stagnated, we carried out a 3D classification (with blush regularization) to separate particles that could result in a high-resolution structure from those of lower quality. The smaller class of 157448 particles, but clearer features, was chosen for the 6^th^ round of 3D refinement (**2.1 Å**) and post-processing (**2.09 Å**). We further performed CTF refinement (CTF with 4^th^ order aberration, anisotropic and parameter fitting), the 7^th^ 3D refinement (**2.05 Å**), post-processing (**2.05 Å**) and polishing. An additional round of 3D refinement (**2.04 Å**), CTF refinement (CTF with 4th order aberration, anisotropic and parameter fitting), the 9^th^ 3D refinement (**2.05 Å**) and post-processing (**2.05 Å**) did not improve the resolution further and we therefore ended the processing in Relion.

### Model building and refinement

The atomic model of *Ficus albivenis* PSI–LHCI (PDB ID: 8WGH) was used as the initial template and rigid-body fitted into the cryo-EM map using UCSF Chimera (Pettersen et al., 2004). The amino acid sequences for all subunits were systematically replaced with the corresponding *Avicennia marina* sequences using Coot (Afonine et al., 2018; Emsley et al., 2010). Pigment cofactors, including chlorophyll a and b, were manually positioned according to the electron density map, with careful validation and adjustment for optimal fit.

For the core subunits PsaA, PsaB, PsaC, and PsaI, publicly available *A. marina* sequences were used for substitution. For subunits lacking sequence information in *A. marina*, initial models were generated automatically using ModelAngelo in sequence-independent mode (without a FASTA input), enabling tracing of the polypeptide backbone and tentative side-chain identification (Jamali et al., 2024). Further refinement and side-chain assignment were performed by comparative analysis of the EM density and sequence conservation, using BLAST alignments within ChimeraX to guide residue placement for highly conserved regions (Meng et al., 2023). All models were iteratively improved and validated by real-space refinement using the Phenix suite, ensuring optimal geometry and fit to the experimental density (Adams et al., 2010).

### Phylogenetic analysis

To compare sequences of light harvesting complex subunits (chains 1-4) with related species, we used sequences embedded in PDB structures and aligned them with the Clustal Omega algorithm in Geneious Prime (**Extended Fig. 5 and 6**). Sequences of all PSI subunits of *Avicennia marina* and related plant species are available in **Extended file 1**.

## Acknowledgement

The cryo-electron microscopy performed in this study was financed by the KAUST Baseline and funds. We are thanking the cryo-EM core lab members at KAUST for their support with the cryo-EM data collections and for maintaining the microscopes and related equipment.

We gratefully acknowledge the Core Technology Platform (CTP) at NYU Abu Dhabi for technical support, and extend particular thanks to Drs. Rachid Rezgui, Mostafa Khair, Sneha Thomas, Liaqat Ali, Muhammad Shiraz Ali, and Lawrence Torres. We also thank the High Performance Computing Center at NYU Abu Dhabi for providing the computational infrastructure and data storage essential for this work. Special appreciation is extended to Dr. Sangram Gore for assistance with field collection of Avicennia marina leaf material. We further thank the Environment Agency – Abu Dhabi and Jubail Mangrove Park for granting permission to collect plant samples for this study.

## Extended figures

**Extended Figure 1:**
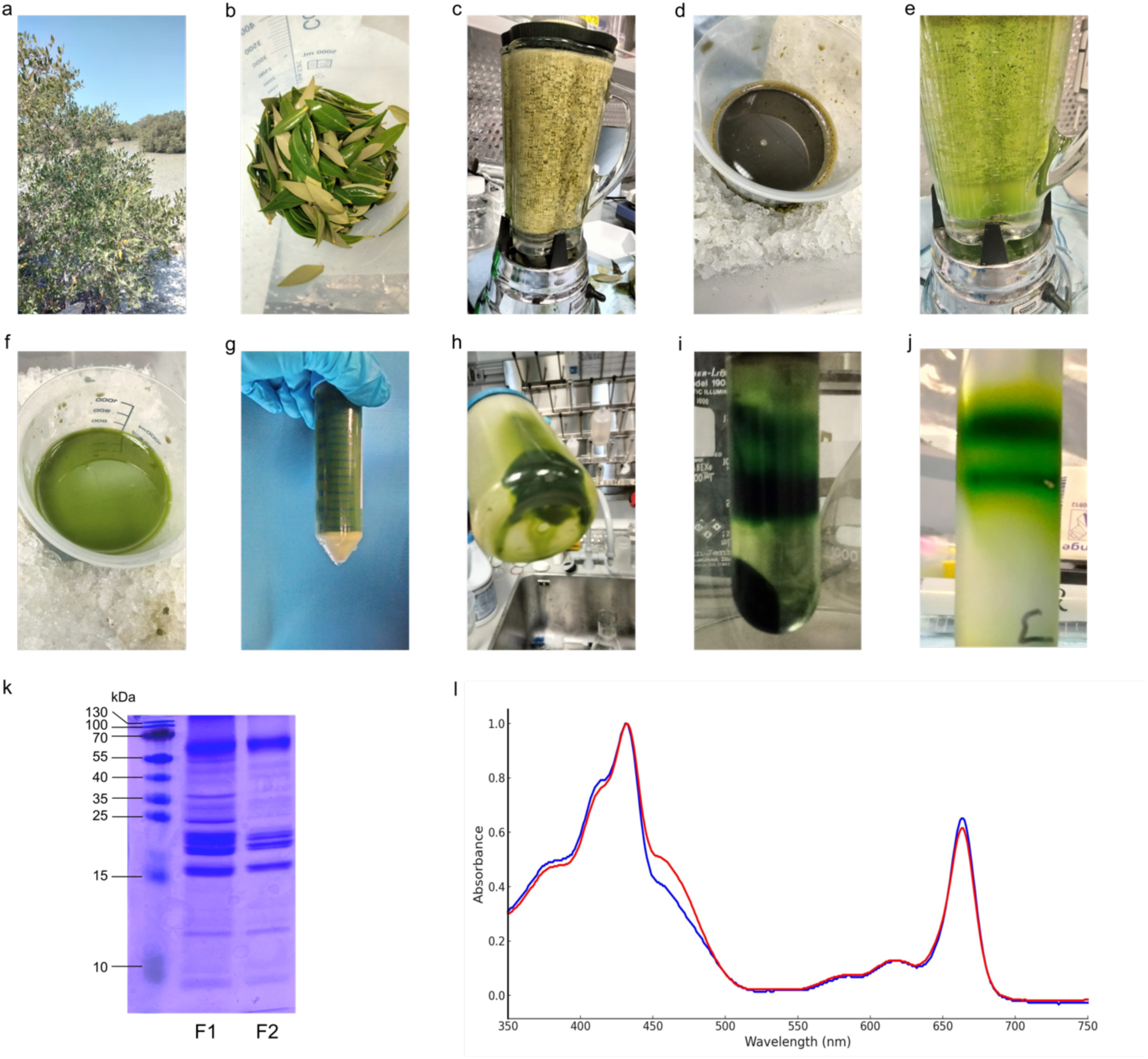
Photosystem I purification from native environment. **a)** Mangrove tree (*A. marina*) in Jubail Mangrove Park, Abu Dhabi. **b)** Freshly harvested leaves were washed thoroughly to remove surface salt excretions. **c–d)** Initial homogenization using conventional plant PSI extraction protocols resulted in oxidation due to high phenolic content, turning the homogenate gray. **e–f)** Modified extraction procedures included phenolic suppressants, yielding a stable green extract. **g–h)** Centrifugation removed starch and insoluble debris from the lysate. **i)** Intact chloroplasts were isolated via Percoll density gradient centrifugation. **j)** Sucrose gradient purification of PSI revealed distinct bands; the lower band corresponding to PSI was collected and concentrated. **k)** SDS–PAGE analysis of both sucrose gradient bands showed higher purity in the lower band, indicative of enriched PSI complex. **l)** Room-temperature absorption spectra of the isolated PSI complex confirmed pigment integrity and spectral characteristics consistent with functional PSI.

**Extended Figure 2:**
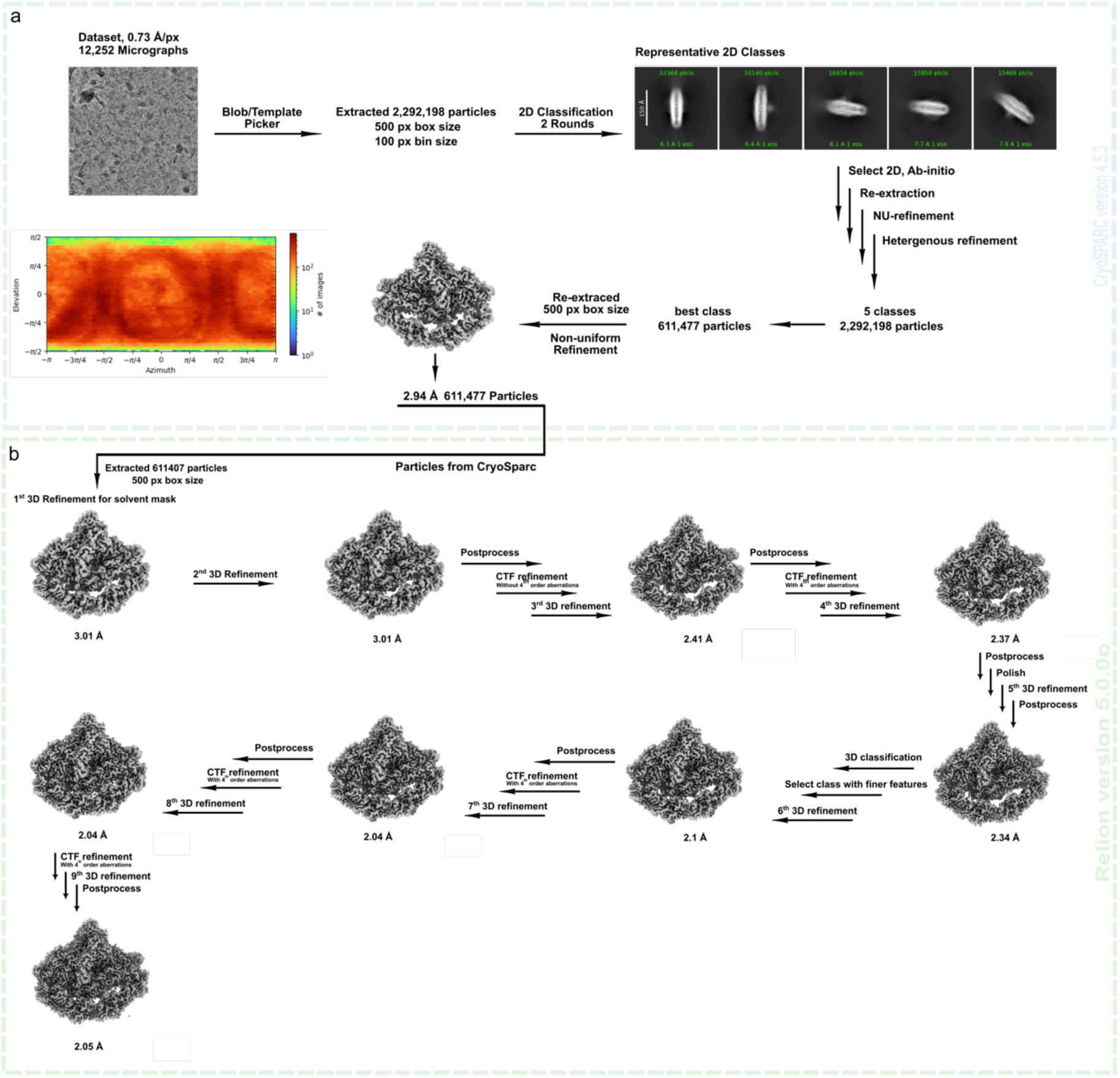
Cryo-electron microscopy processing workflow. **a)** CryoSparc workflow: A dataset of 12,252 micrographs at a resolution of 0.73 Å /px was collected. From these micrographs, 2,292,198 particles were extracted, binned to 100 px box size and 2D classified. Selected 2D classes were used for ab initio, re-extracting particles at 500 px, non-uniform refinement, heterogenous refinement, 3D classification and again non-uniform refinement. 611,477 particles were exported into Relion5. **b)** Relion5 workflow. Re-iterative rounds of refinement, postprocess, 3D classification, CTF refinement and polishing resulted in a structure with an overall resolution of 2.05 Å.

**Extended Figure 3:**
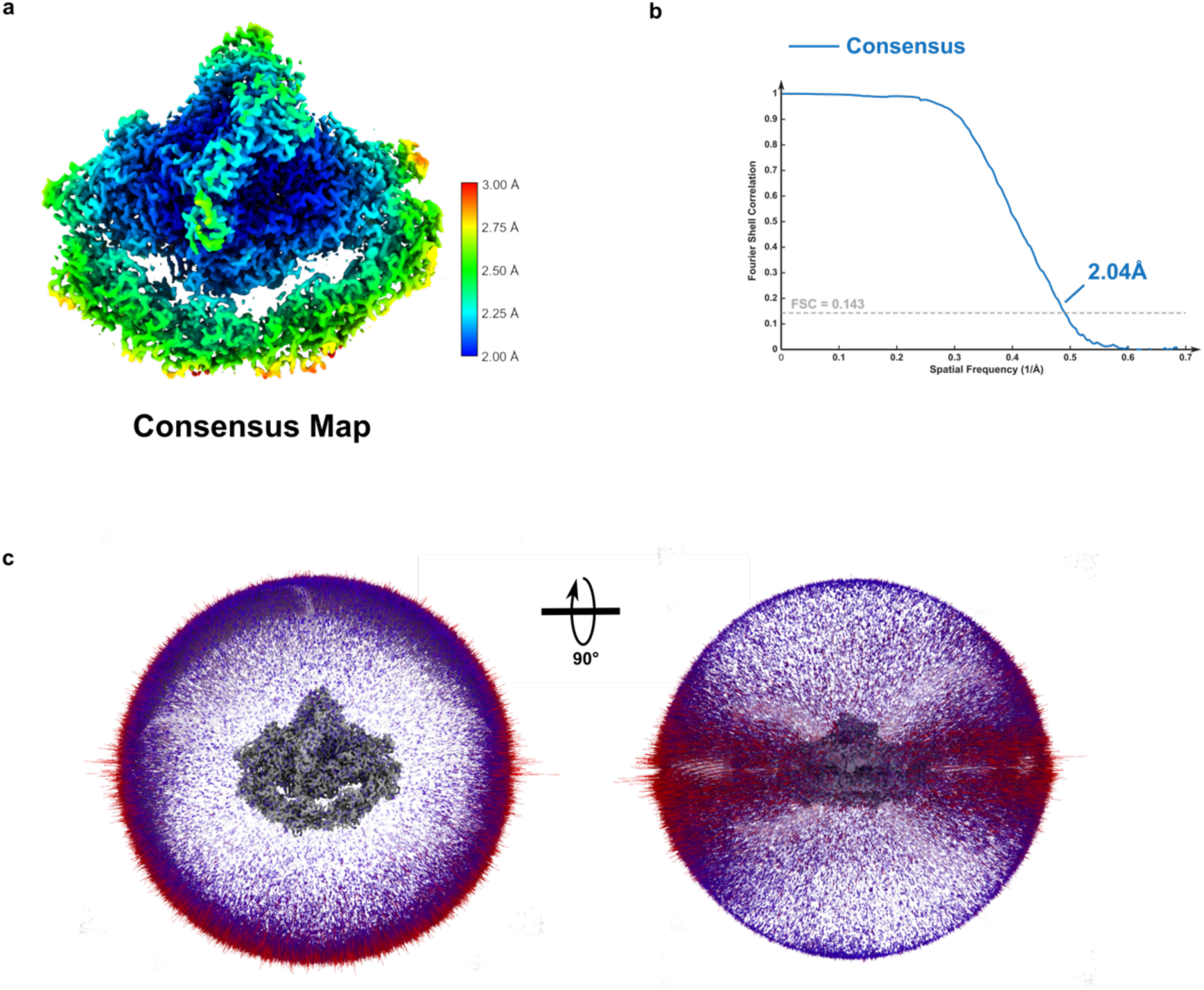
Cryo-electron microscopy processing workflow quality control. **a)** Local resolution of consensus map. **b)** Fourier shell correlation (FSC) curves with gold standard line (FSC = 0.143). **c)** Angular distribution of consensus map.

**Extended Figure 4:**
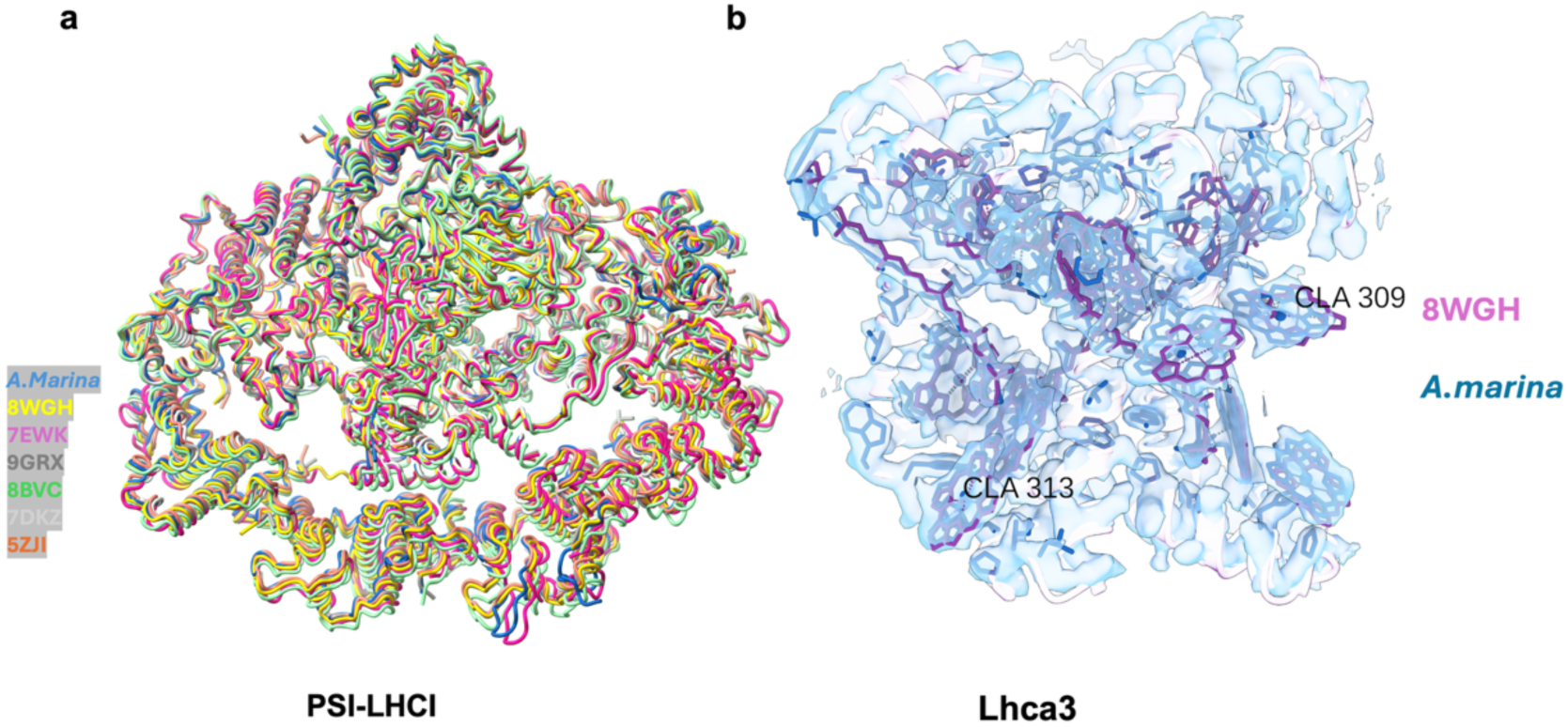
Structural comparison of *A. marina* PSI–LHCI with other plant PSI–LHCI supercomplexes. **a)** Superimposed stromal views of PSI–LHCI protein backbones, shown in cartoon representation and colored as follows: *A. marina* (blue), 8WGH (yellow), 7EWK (magenta), 9GRX (dark), 8BVC (green), 7DKZ (light grey), and 5ZJI (orange). Cofactors are omitted for clarity, highlighting the conserved architecture of the PSI core and LHCI belt across species. **b)** Cryo-EM density of the Lhca3 subunit in *A. marina*, overlaid with modeled cofactors. Three regions in Lhca3 show notable divergence in the vicinity of a low-energy chlorophyll dimer, as compared to *F. albivenis* (PDB: 8WGH, magenta). The dimeric red chlorophyll positions are well defined in the density, with the primary sites corresponding to 309/310 and 303/313, reflecting subtle shifts relative to reference structures.

**Extended Figure 5:**
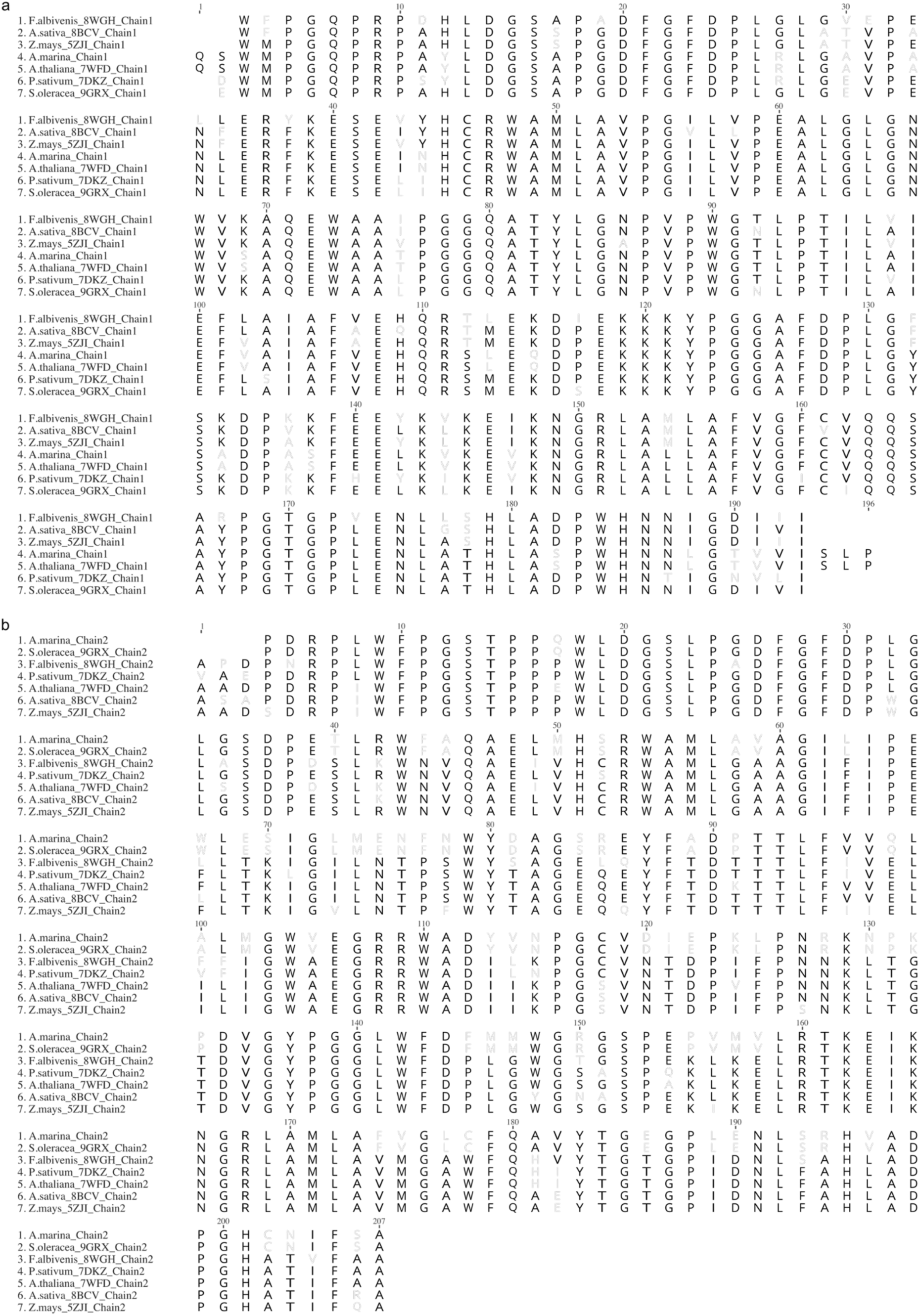
Sequence alignments of related plant species. **a)** Chain 1 of *Avicennia marina* PSI aligned with chain 1 of *A. sativa*, *A. thaliana*, *F. albivenis*, *P. sativum*, *S. oleracea*, *Z. mays*. **b)** Chain 1 of *Avicennia marina* PSI aligned with chain 1 of *A. sativa*, *A. thaliana*, *F. albivenis*, *P. sativum*, *S. oleracea*, *Z. mays*.

**Extended Figure 6:**
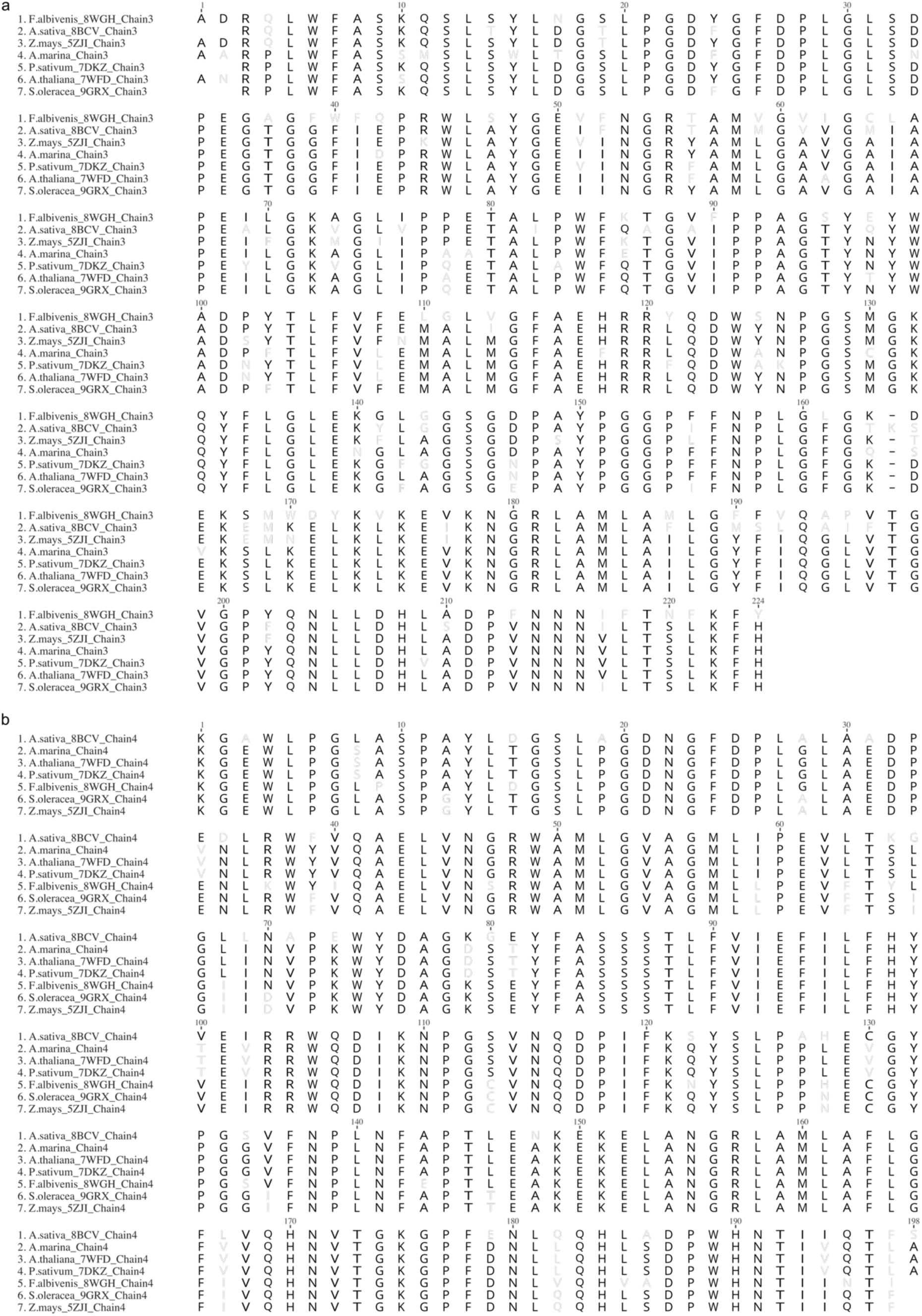
Sequence alignments of related plant species. **a)** Chain 3 of *A. marina* PSI aligned with chain 1 of *A. sativa*, *A. thaliana*, *F. albivenis*, *P. sativum*, *S. oleracea*, *Z. mays*. **b)** Chain 4 of *Avicennia marina* PSI aligned with chain 1 of *A. sativa*, *A. thaliana*, *F. albivenis*, *P. sativum*, *S. oleracea*, *Z. mays*.

## Notes

### Competing Interest Statement

The authors have declared no competing interest.

### Summary of Updates

Labels of Fig 3 and 4 are changed.

